# Genome-microbiome interplay provides insight into the determinants of the human blood metabolome

**DOI:** 10.1101/2022.02.04.479172

**Authors:** Christian Diener, Chengzhen L. Dai, Tomasz Wilmanski, Priyanka Baloni, Brett Smith, Noa Rappaport, Leroy Hood, Andrew T. Magis, Sean M. Gibbons

**Author notes:** equal contribution (Christian Diener, Chengzhen L. Dai).

## Abstract

Variation in the blood metabolome is intimately related to human health. Prior work has shown that host genetics and gut microbiome composition, combined, explain sizable, but orthogonal, components of the overall variance in blood metabolomic profiles. However, few details are known about the interplay between genetics and the microbiome in explaining variation on a metabolite-by-metabolite level. Here, we performed analyses of variance for each of the 945 blood metabolites that were robustly detected across a cohort of 2,049 individuals, while controlling for a number of relevant covariates, like sex, age, and genetic ancestry. Over 60% of the detected blood metabolites were significantly associated with either host genetics or the gut microbiome, with more than half of these associations driven solely by the microbiome and around 30% under hybrid genetic-microbiome control. The variances explained by genetics and the microbiome for each metabolite were indeed largely additive, although subtle, but significant, non-additivity was detected. We found that interaction effects, where a metabolite-microbe association was specific to a particular genetic background, were quite common, albeit with modest effect sizes. The outputs of our integrated genetic-microbiome regression models provide novel biological insights into the processes governing the composition of the blood metabolome. For example, we found that unconjugated secondary bile acids were solely associated with the microbiome, while their conjugated forms were under strong host genetic control. Overall, our results reveal which components of the blood metabolome are under strong genetic control, which are more dependent on gut microbiome composition, and which are dependent upon both. This knowledge will help to guide targeted interventions designed to alter the composition of the blood metabolome.

## Introduction

The human blood metabolome is shaped by a combination of intrinsic and extrinsic forces and constitutes the primary resource pool for human metabolism. While the composition of the plasma metabolite pool is strongly driven by diet, lifestyle, and the ecology of the gut microbiota, the fate of individual metabolites in the blood is often tightly regulated by host genetics ^1^.

Genetic variants are known to alter the human blood metabolome in several disease-relevant contexts. For example, certain deleterious alleles impact cholesterol levels and contribute to hypercholesterolemia, while another allele drives phenylalanine accumulation and leads to phenylketonuria ^2–4^. The majority of genetic variants associated with blood metabolite levels affect either enzymes or solute carriers, thus directly influencing an individual’s ability to produce, consume, secrete, or absorb small molecules ^5,6^. Many of the currently identified genetic variants are pleiotropic ^7^, which indicates a vast interconnectedness of the metabolome to different systems of the body.

Recent studies have identified the human gut microbiome as a major determinant of blood metabolite variability ^1,8^. In this prior work, host genetics and gut microbiome composition were found to associate with plasma metabolite levels in a largely orthogonal manner, which is supported by the observation that, for the most part, host genetics and the gut microbiota do not tend to associate strongly with one another ^9^, even though a number of clear examples of genome-microbiome associations have been shown ^10,11^.

Despite these broad, multivariate regression results showing a global correspondence between genomics, the microbiome, and the metabolome, little is known about how this maps onto individual blood metabolites. While one might expect that the variation in metabolites produced by bacteria in the human gut is mostly governed by the microbiota and that metabolites specific to human metabolic processes are associated more with host genetics, there are examples, as in the case of microbe-host co-metabolites like conjugated secondary bile acids ^12^, where the story becomes more complicated.

Intestinal signaling, such as the activation of FXR and TGR5 receptors in the human gut, can regulate glucose, insulin, cholesterol, and bile acid homeostasis ^13–16^. Furthermore, many microbiome-derived metabolites are modified by hepatic enzymes and converted into a variety of conjugated compounds, such as hippurate or polyamines. These multi-layer filters on microbe-host co-metabolism make it challenging to map blood metabolites to potential microbial precursors ^17–20^. For example, blood cholesterol levels are affected by host genetic variants, but they can also be regulated through intestinal signaling, involving the production of secondary bile acids by gut commensals, or through gut microbial cholesterol dehydrogenases that funnel host cholesterol into fecal coprostanol ^21–23^. Thus, even though host genetics and the microbiome appear to have largely orthogonal effects on the blood metabolome as a whole, they nonetheless act on an overlapping set of metabolites, potentially explaining independent components of the variance of these compounds.

Additionally, even in the absence of strong heritability of human gut commensals, there are key examples where genetics and the microbiome may interact in particular disease conditions, such as cystic fibrosis ^24^. This raises the question of whether there are instances where microbiome-metabolome associations are modulated by the genetic background of the host in a similar manner as other gene-environment interactions ^25^, where a genetic variant may augment or attenuate an individual’s risk of developing a particular disease when exposed to a specific environmental risk factor.

Here, we studied variability in plasma metabolite abundances in a cohort of 2,049 healthy individuals from the Pacific West of the US. We find extensive interplay between the genetic and gut microbial determinants of individual metabolite levels in the blood, which provides deep insights into the microbe-host co-metabolisms that govern the composition of the human blood metabolome.

## Results

### Identifying plasma metabolites associated with genetic features only, gut microbiome features only, or with both genetic and microbial features

To identify associations between host genetics and individual circulating blood metabolite levels, we performed genome-wide association analyses on 7.68 million common variants (Minor Allele Frequency ≥ 1%) in 2,049 individuals for each of the 945 detected blood metabolites. A total of 299 metabolites of the 945 tested (31.6%) were associated with one or more of the 389 independent lead variants that passed the genome-wide significance threshold (p < 5.29 × 10^−11^; see Methods and Fig. S1). Of these 389 variants, 123 were in intergenic regions and 266 variants mapped to 166 genes, including those associated with inborn errors of metabolism such as *ACADS*, *MTHFR*, *DMGDH*, and *ETFDH*. Potential loss-of-function consequences were predicted for three variants: stop gained for rs183603441 in *HYKK*, stop lost for rs358231 in *GBA3*, and a splice donor variant for rs114286107 in *AGXT2*. Missense mutations were predicted for 40 variants, affecting 74 metabolites. However, the majority of the metabolite-associated variants found within genes were synonymous.

Comparison with associations from previous metabolomics studies and the GWAS catalog revealed 6 novel genomic loci that influence metabolite levels. These novel metabolic quantitative trait loci (mQTL) were associated with 7 metabolites. In particular, lactosylceramide lactosyl-N-nervonoyl-sphingosine (d18:1/24:1) was associated with lead variants rs3752246 (P = 8.5 × 10^−26^) located in *ABCA7* and rs12979724 (P = 8.5 × 10^−12^) located near *CNN2*. rs3752246 is a missense variant and has been previously identified as a risk variant for late-onset Alzheimer’s disease ^26–28^. To test whether these two traits share the same causal SNP at this genomic locus, we performed colocalization analysis and found the posterior probability of causal SNP sharing to be 99.6% (see Materials and Methods). Prior studies have identified altered levels of ceramides in the blood and brain of individuals with Alzheimer’s Disease, with the d18:1/24:1 ceramide showing a strong and robust association with Alzheimer’s Disease in a recent meta-analysis ^29–31^. Our results suggest a possible shared genetic architecture underlying lactosyl-N-nervonoyl-sphingosine (d18:1/24:1) levels and late-onset Alzheimer’s disease.

Associations with more than one metabolite were identified for 82 of the 389 significant lead variants (21.1%), revealing the extent of pleiotropy. The effect sizes of pleiotropic variants are generally higher than non-pleiotropic variants (P = 3.67 × 10^−3^; two-sided Mann-Whitney U-test), while the distributions of minor allele frequency were similar between the two types of variants. Overall, the 82 pleiotropic lead variants were associated with 186 metabolites (62.2% of all significant genetically-associated metabolites). Four variants (rs4149056, rs1047891, rs148982377, and rs11568824), of which rs4149056 and rs1047891 are missense variants, were each associated with more than 10 metabolites. rs4149056 in *SLCO1B1* (solute transporter in liver) was associated with 19 metabolites, including primary and secondary bile acids, conjugates of polyunsaturated fatty acids, and free fatty acids. rs1047891 in *CPS1* (mitochondrial enzyme involved in the urea cycle) was found to be associated with 11 metabolites, many of which were conjugated to glycine and glutamine moieties. rs148982377 in *ZNF789* and rs11568824 in *ZSCAN25* (transcription factors) were associated with the same set of 11 steroid hormone metabolites, primarily conjugates of DHEA and androsterone.

Analysis at the gene level reveals a further degree of pleiotropy as well as polygenicity. Using MAGMA ^32^, we identified associations between genes and metabolites, accounting for linkage disequilibrium. 242 metabolites were associated with at least one gene, with 351 significant genes identified in total (P < 9.48 × 10^−9^). Of these, 128 genes (36.47%) were associated with more than one metabolite. Individual pleiotropic genes tend to be associated with metabolites with similar biochemical properties, providing insights into unidentified metabolites. For example, the gene cluster of *UGT1A1*, *UGT1A3*, *UGT1A4*, *UGT1A5*, *UGT1A6*, *UGT1A7*, *UGT1A8*, *UGT1A9*, and *UGT1A10* was associated with 13 metabolites, including bilirubin, biliverdin, and eight unidentified compounds. These genes encode UDP-glucuronosyltransferase (UGT) enzymes, which metabolize bilirubin ^33,34^. It was later determined by Metabolon that the eight unidentified compounds were degradation products of bilirubin. We also observe instances of polygenic associations among many metabolites. For example, five of the 53 metabolites associated with the fatty acid desaturase (FADS) gene cluster were also associated with the hepatic lipase gene *LIPC*. We also observed shared genetic architecture for other combinations of blood metabolites and complex traits. For example, we found associations between the missense variant rs1260326 in *GCKR* with mannose (P = 1.2 × 10^−44^) and 1-carboxyethyl-valine (P = 2.9 × 10^−11^). rs1260326 has been identified as a risk variant in a genome-wide association meta-analysis of type 2 diabetes and Crohn’s disease ^35^. Colocalization analysis of these four traits identified sharing of a causal SNP at this locus (with posterior probability > 0.9).

In order to identify associations between the gut microbiome and circulating blood metabolite levels, we performed regressions using centered-log-ratio (CLR) transformed bacterial genus-level abundances as independent variables, while correcting for sex, age, sex-age interactions, BMI, and genetic kinship (i.e., the first 5 principal components of the genetic distance matrix). A majority of the tested metabolites had significant associations with at least one bacterial genus in the gut microbiome (522/945 = 55.2%, with FDR-corrected p<0.05). Here, the average fraction of explained variance (R^2^) was slightly lower when compared to genetic associations (mean R^2^ of 0.04 and 0.09 for microbiome and host genetic features, respectively).

The blood metabolites with the largest fraction of variance explained by microbial features were dominated by compounds involved in bacterial metabolism of aromatic and phenolic compounds, such as cinnamoylglycine, hydrocinnamate, hippurate, and phenylacetylglutamine (all R^2^>0.2, Fig. 1D). Catabolism of phenylalanine and phenylacetate to cinnamate and benzoate is exclusive to the microbiome and is conspicuously absent in germ-free mice ^36,37^. Hippurate is formed in the liver by conjugating benzoate to a glycine and blood levels of hippurate have been positively associated with gut microbiome alpha-diversity ^8,38^ and with overall metabolic health ^39^.

**Figure 1.**
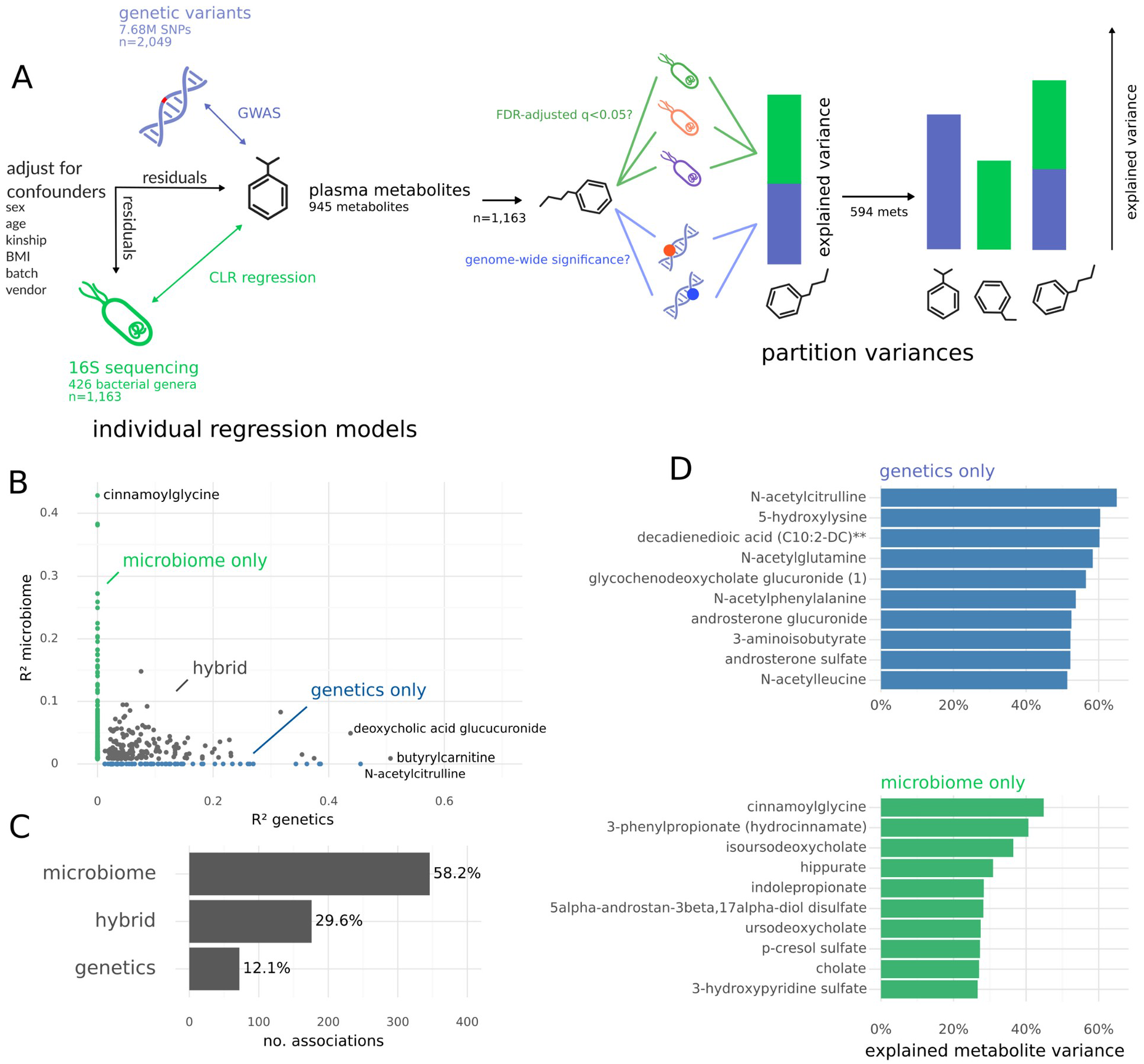
Study design and cross-sectional variances in metabolites explained by host genetics, gut bacterial genera, or both. (A) The study comprised a metabolome-genome-wide association analysis and a metabolome-microbiome-wide association analysis (regressions on centered log-ratio transformed bacterial genus-level abundances) performed in the Arivale cohort. Explained variance for a specific blood metabolite can be partitioned into host genetic and microbiome associated components. (B) Fraction of variance (R^2^) explained by host genetic or microbiome features across the 594 metabolites with significant associations with either the genome or the microbiome (FDR-corrected p<0.05). Specific sub-groups are annotated in the plot, including metabolites only associated with host genetics, metabolites only associated with gut genera, or metabolites significantly associated with both (hybrid). (C) Percentage of the 594 significant metabolites associated only with host genetics, only with the microbiome, or with both (hybrid). (D) The 10 metabolites with the highest explained variance by either genetics only (blue) or by the microbiome only (green).

Additionally, we found several products of bacterial protein fermentation, such as p-cresol derivatives and the tryptophan breakdown product indolepropionate, associated with the gut microbiome (Fig. 1D). Furthermore, we found that all secondary bile acids identified in plasma were significantly associated with the gut microbiome (FDR-corrected p<0.05), which is expected because primarily bile acids are deconjugated into secondary bile acids by bile salt hydrolases expressed by gut commensal bacteria ^40^.

### Genetic and microbial factors additively contribute to explained variance in blood metabolite abundances

We identified a total of 594 out of 945 plasma metabolites (62.7%) that were associated with either genetic factors, microbial factors, or both (Fig. 1A-B). Most of these metabolites (522) were significantly associated with the microbiome. 29.6% of these metabolites showed “hybrid” associations, meaning they were associated with genetic as well as microbial factors (Fig. 1A-B). In particular, we found that about ¾ of all metabolites with a significant genetic association also showed a microbial association (i.e., 176 of 248 metabolites associated with genetic variants). Conversely, only ⅓ of all metabolites with a microbiome association also showed a hybrid association with host genetics. Consistent with the average explained variances of metabolites reported above for genetics and the microbiome, hybrid metabolites with particularly large explained variances (i.e., >20%) showed a tendency to have higher genetic R^2^ values than microbial R^2^ values (see Fig. 1B and Fig. 2A).

**Figure 2.**
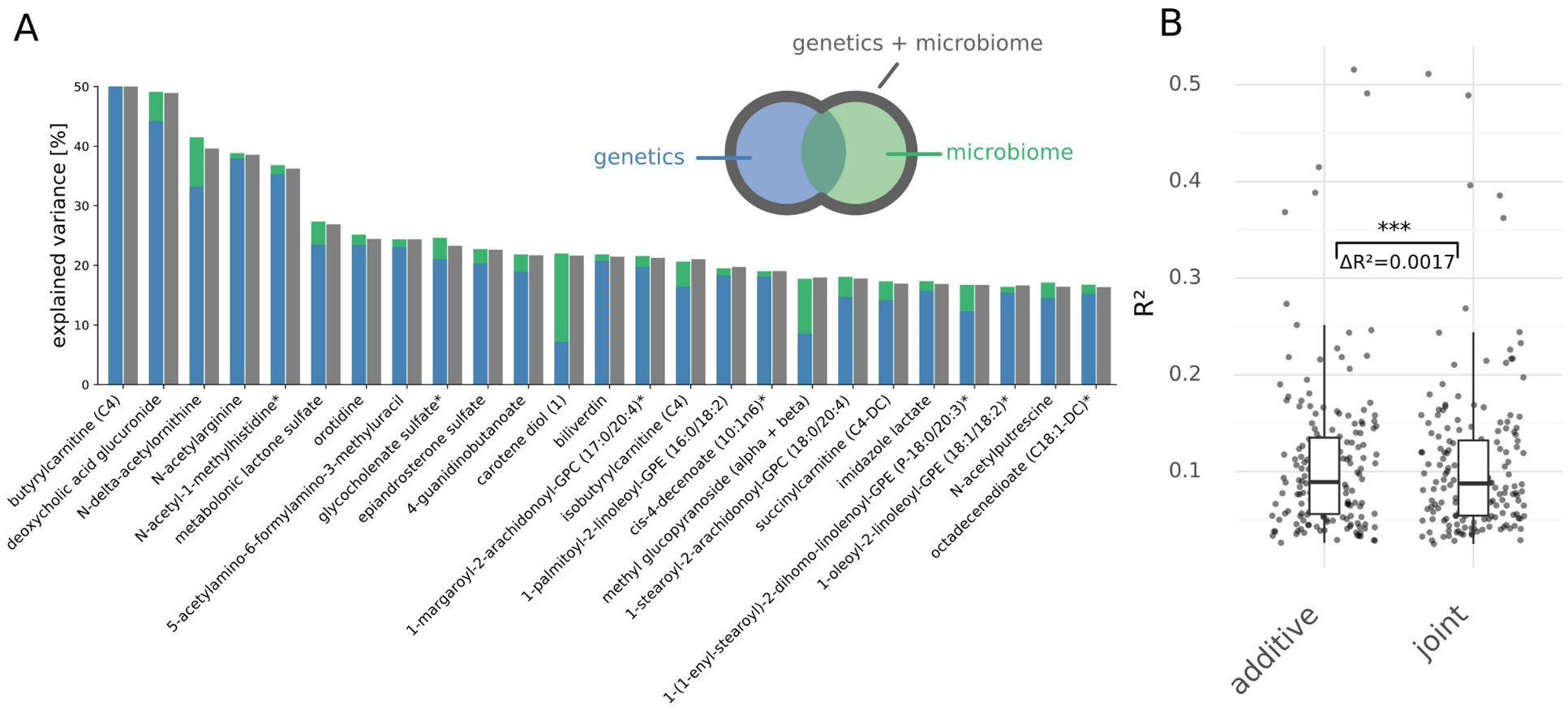
Metabolites associated with both genetics and the microbiome. (A) The 20 metabolites with the highest total R^2^ that were significantly associated with both host genetics and the microbiome. Blue and green colored bars denote individual R^2^ values in genetics-only and microbiome-only regression models, respectively. Gray bars denote R^2^ values from regression using joined genetics and microbiome data. 0.9% of the variance in butyrylcarnitine plasma abundance was explained by microbial features (i.e. too little to be visible in the barplot). (B) R^2^ values obtained by either adding individual contributions of genetics and the microbiome (additive) or by performing a joint regression (joint). The difference between the two groups indicates a small, but nonetheless significant, overlap in variance explained by genetics and the microbiome. However, the variances explained by host genetics and the microbiome were largely additive. Stars denote significance (*** - p<0.001).

In order to test whether genetic and microbial factors contained redundant information, we compared a joint genetics-microbiome model to individual genetic and microbial models. If the overlap in variance explained between genetic and microbial feature sets was large, the joint model R^2^ value would be substantially smaller than the sum of the individual model R^2^ values (see Fig. 2A). Although we found that the difference in R^2^ between the joint model and the sum of the individual model R^2^ values was statistically significant across a very large number of models, the magnitude of this difference was extremely small (median difference in R^2^<0.0014), indicating that genetics and the microbiome explain nearly exclusive components of the variance for a given metabolite. This result is consistent with prior work showing that the variances in the blood metabolome explained by the microbiome and host genetics were largely orthogonal ^1,9^. However, here we demonstrate that this is not only true globally, but at the individual metabolite level as well.

Finally, we investigated whether the genetic background of the host could modulate microbiome-metabolite associations (see Fig. 3A). To this end, we tested all genetic variant / bacterial genus / metabolite triplets, filtering for only those genetic variants or genera that were previously associated with at least one metabolite (i.e., 16,905 triplets in total). We employed a strategy similar to gene-environment interaction studies, using the CLR transformed abundances of microbial genera as the environmental variable, while adjusting for covariates (see Methods) ^41^. 207 interaction effects were deemed significant under an FDR-corrected p-value cutoff of 0.05, involving 49 distinct metabolites. Though gene-microbiome interactions were quite common, they only explained very small fractions of plasma metabolite variability (mean R^2^=0.005). The amount of explained variance by the interaction terms tended to correlate negatively with variance explained by the genetic variant itself (Fig. 3B, Pearson’s rho=−0.43, p=6.8e-11).

**Figure 3.**
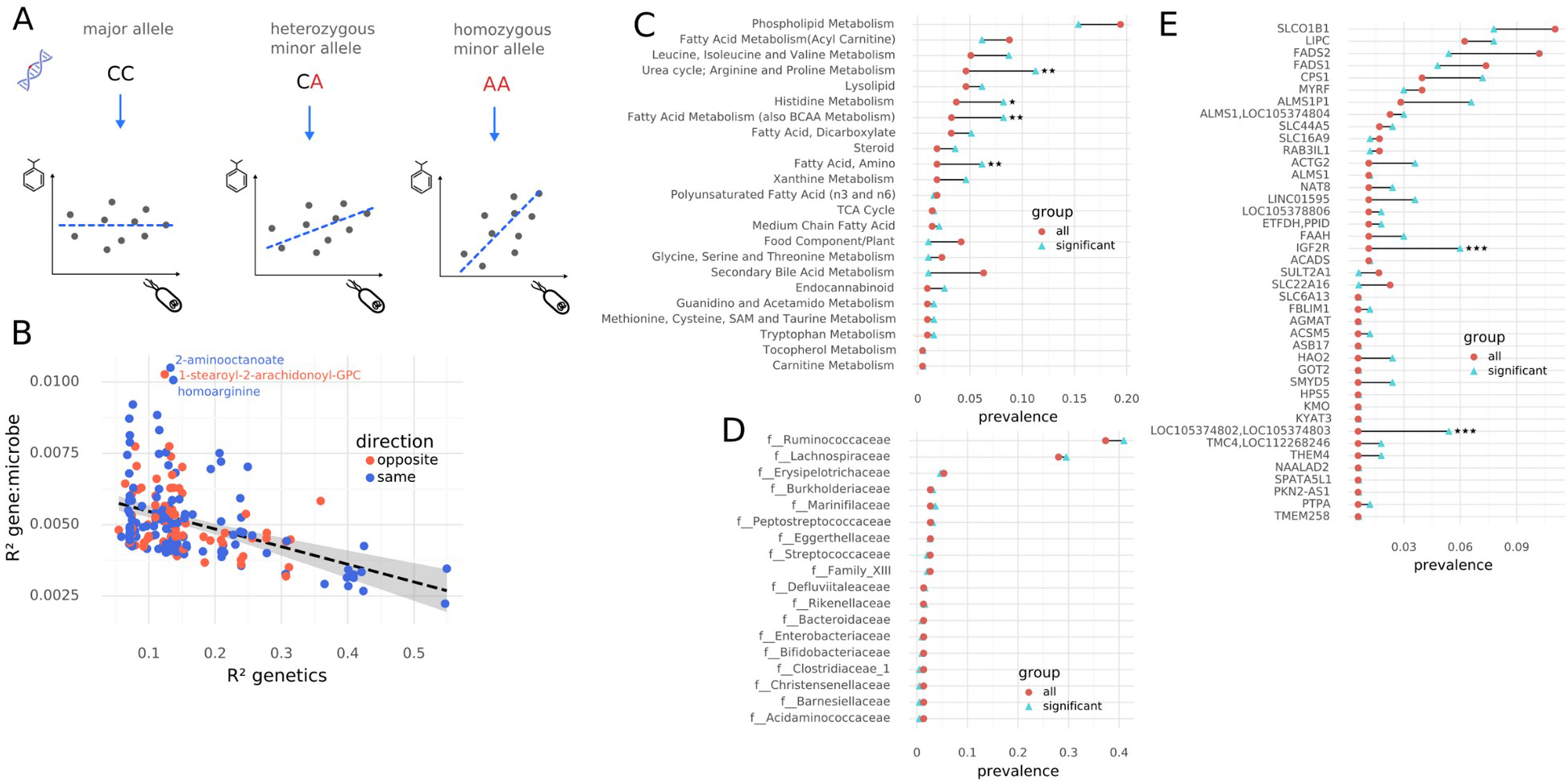
Gene-microbiome interactions. (A) Gene-microbiome interactions occur when the correlation between a gut bacterial genus and a blood metabolite is itself conditional on a specific allele. (B) R^2^ values for genetic associations and for the corresponding gene-microbe interactions. Each dot denotes a single model containing both terms. Interaction terms and genetics-only terms are negatively correlated (rho=-0.43, p=6.8e-11). (C) Pathway enrichment analysis for metabolites with significant gene-microbe interactions. (D) Bacterial family-level enrichment analysis for significant gene-microbe interactions. (E) Host gene enrichment analysis for significant gene-microbe interactions. In (C-E) red circles denote prevalence in all tested features (i.e., background prevalence) and blue triangles denote prevalence in only those features with significant interaction terms. Stars denote significantly enriched features under a one-sided hypergeometric test with Benjamini-Hochberg correction for multiple testing (* - q<0.05, ** - q<0.01, *** - q<0.001).

Gene-microbiome interactions were significantly enriched in metabolites involved in the urea cycle, histidine metabolism, and fatty acid metabolism (Fig. 3C, one-sided hypergeometric test, FDR-corrected p<0.05) and also showed enrichment in the *IG2F* gene (insulin-like growth factor 2) and two overlapping loci on chromosome 2 (Fig. 3E, one-sided hypergeometric test FDR-corrected p<0.05). No enrichments in bacterial phyla, genera, or families were observed (Fig. 3D). The highest fraction of explained variance for these interaction terms was observed for a handful of distinct triplets, including the metabolites homoarginine, 2-aminooctanoate, and 1-stearoyl-2-arachidonoyl-GPC. Low levels of homoarginine are associated with a higher risk of cardiovascular disease ^42,43^ and we observed a negative gene-microbe interaction between the missense variant rs1047891 in the *CPS1* gene (urea cycle) and several genera in the *Ruminococcaceae* family, where the genus abundance tended to show a negative correlation with the plasma homoarginine abundance only in heterozygous individuals carrying the C→A minor allele (Fig. 4).

**Figure 4.**
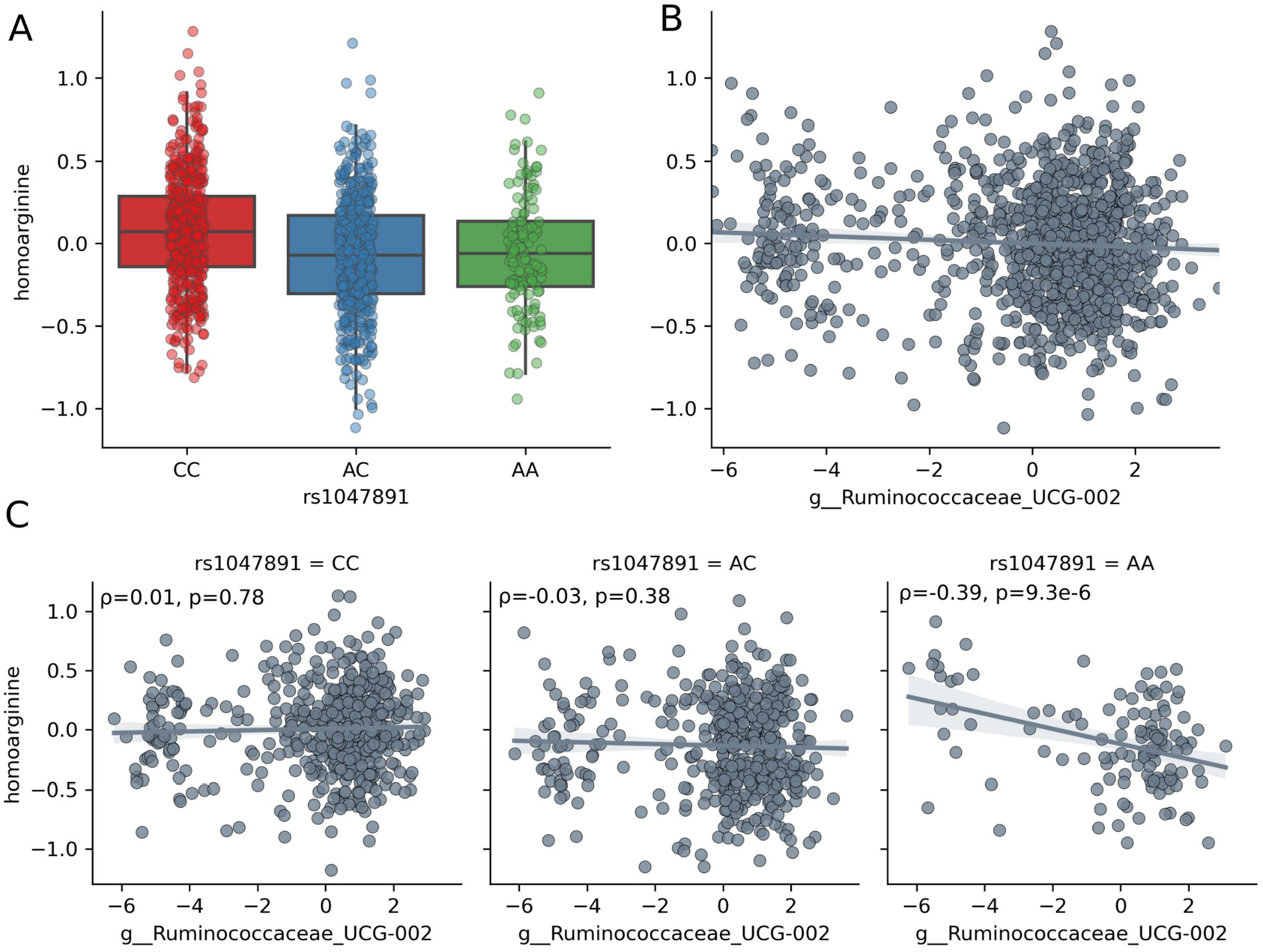
Gene-microbiome interactions in explaining variation in homoarginine levels. (A) Homoarginine levels across the alleles of rs10447891 locus located in the *CPS1* gene. (B) Homoarginine levels plotted against centered log-ratio transformed genus-level abundances of *Ruminicoccaceae UCG-002* across all host genotypes. (C) Homoarginine levels against centered log-ratio transformed genus-level abundances of *Ruminicoccaceae UCG-002* stratified by genotype. In (A-C) homoarginine abundances are log-transformed abundances adjusted for common confounders, as described in Materials and Methods. In (B-C) “ρ” denotes the Pearson Product-Moment correlation coefficient of the regression and “p” the p-value under a Pearson correlation test. The solid line indicates the linear regression line and the shaded area is the 95% confidence interval of the regression. Associations were corrected for sex, age, age^2^, sex:age, and sex:age^2^ interactions, BMI, microbiome vendor, metabolomics batch, and the first 5 principal components of genetic ancestry.

### Genetics strongly associated with the fate of conjugated, but not unconjugated, secondary bile acids

Having mapped plasma metabolite variability to genetic and microbial factors, we now asked whether the observed partitioning of variances explained may change along pathways that involve host-microbe co-metabolisms. To this end, we investigated the metabolism of secondary bile acids. Secondary bile acids are formed in the large intestine via microbial deconjugation of primary bile acids, reabsorbed into the bloodstream via the portal vein, and further metabolized in the liver ^44^. Thus, secondary bile acid levels in the bloodstream are influenced by both the microbiome and the host. We investigated whether individual bile acid species variances were predominantly explained by genetic factors, microbial factors, or both.

While unconjugated secondary bile acids were exclusively associated with the gut microbiome, 5 out of 6 detected secondary bile acid conjugates fell into the hybrid class (Fig. 5A). In particular, plasma deoxycholate abundances showed no significant genetic contribution, whereas more than 40% of the variance in the conjugates deoxycholate glucuronide and deoxycholate sulfate was explained by host genetics (Fig. 5B). Other glucuronidated non-bile-acid compounds, such as p-cresol glucuronide, did not show this pattern (see Fig. 1D). Modifications like glucuronidation or sulfation usually occur in the liver and are used to mark metabolites for excretion in feces or urine ^45,46^. These results suggest that the clearance of conjugated secondary bile acids from the body is under strong genetic control.

**Figure 5.**
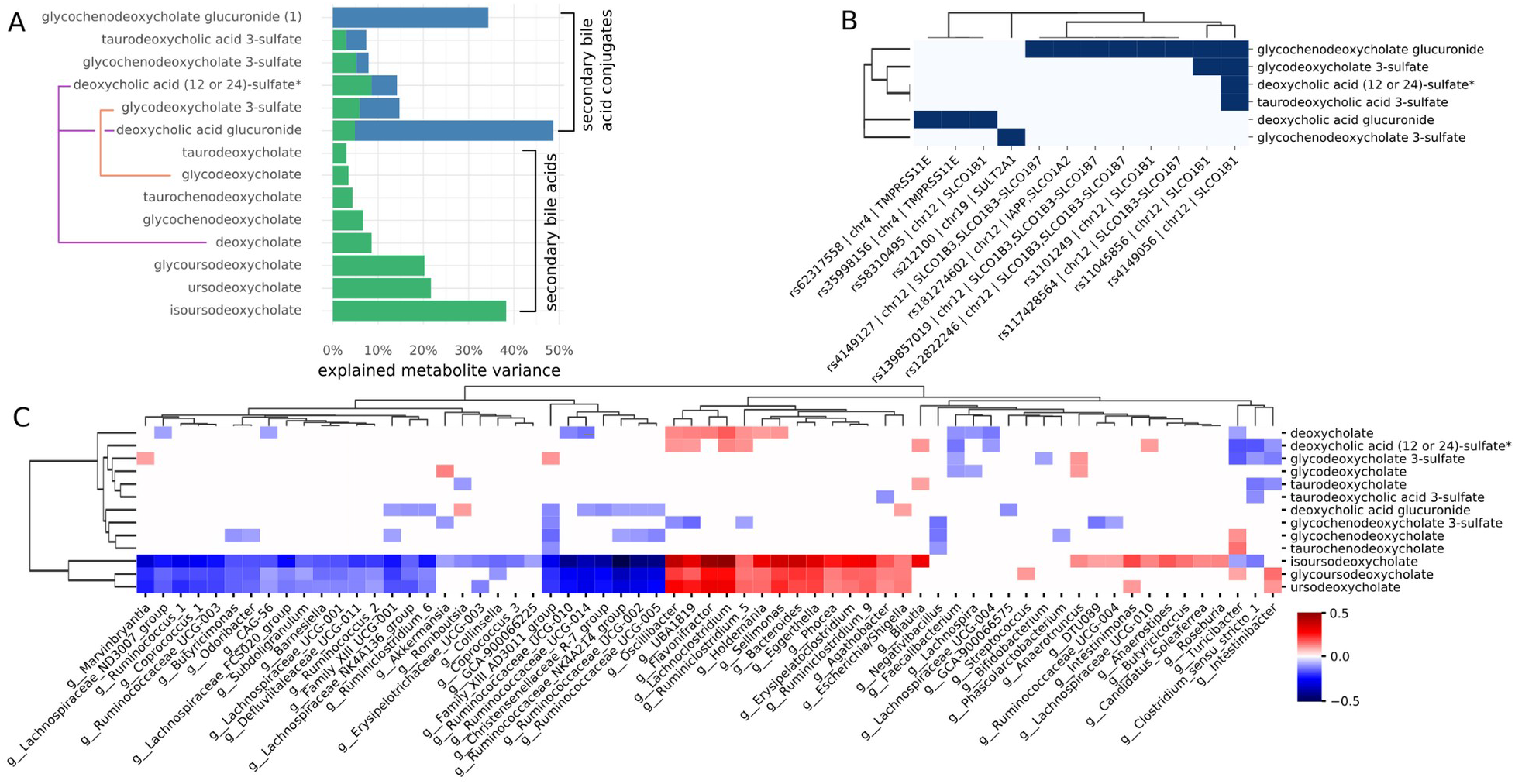
Host genetic and gut microbial associations with secondary bile acids. (A) Explained variances for unconjugated secondary bile acids and hepatic secondary bile acid conjugates. Purple and orange lines denote modifications of the same secondary bile acid. (B) Genetic variants associated with secondary bile acid conjugates. Dark blue cells denote associations that passed a genome-wide significance threshold in the GWAS. (C) Associations between bacterial genera and secondary bile acids (both conjugated and unconjugated). Only significant associations are shown (FDR-corrected and confounder-adjusted q<0.05). Fill colors denote the correlation coefficients (see legend). Associations were corrected for sex, age, age^2^, sex:age, and sex:age^2^ interactions, BMI, microbiome vendor, metabolomics batch, and the first 5 principal components of genetic ancestry. * indicates compounds for which a standard is not available, but Metabolon is confident in its identity; ** indicates a compound for which a standard is not available, but Metabolon is confident in its identity.

All but one hepatically modified secondary bile acid showed associations with both genetic and microbial features. The one exception was glycochenodeoxycholate glucuronide, which was only associated with genetics and not with the microbiome, even though the unconjugated form, glycochenodeoxycholate, was only associated with the microbiome (Fig. 5A). The genomic variant rs4149056 was associated with 4 of the 6 modified secondary bile acids and has been shown previously to affect the abundance of bile acids in plasma ^47^ (see Fig. 5B). This variant is located in the solute carrier protein SLCO1B1, which is expressed in the liver ^48^ and transports secondary bile acids, with a preference for sulfated bile acids and bile salts ^49^. Several other variants in SLCO1B1 were associated with secondary bile acid derivatives. We also identified several variants in the *SLCO1B3-SLC1B7* gene cluster that mostly affected glycochenodeoxycholate glucuronide plasma abundances, and we found several variants located on chromosome 4, one in the TMPRSS11E serine protease and several variants without an associated gene identity, that specifically affected deoxycholate glucuronide levels (Fig. 5B).

Thus, whereas cross-sectional variation in unmodified secondary bile acid levels were not explained by host genetic variation, plasma abundances of several secondary bile acid conjugates were significantly associated with a diverse set of compound-specific genetic variants on chromosomes 4 and 12. Additionally, we observed that association patterns with bacterial genera changed when comparing unmodified deoxycholate to deoxycholate glucuronide (Fig. 5C). Deoxycholate showed mostly positive associations, for instance with *Bacteroides*, *Phocea*, and *Lachnoclostridium*, whereas deoxycholate glucuronide showed mostly negative associations with several genera in the *Ruminicoccaceae* family (Fig. 5C).

### Sphingosines and ceramides show a range of genetic, microbiome, and hybrid associations

We next asked whether hybrid associations would also be common in another disease-relevant class of blood metabolites: ceramides. High ceramide levels have been shown to be associated with insulin resistance, hypercholesterolemia, liver steatosis, and the formation of lipid rafts ^50–54^. Furthermore, some classes of ceramides in the blood increase the risk for late-onset Alzheimer’s disease, as they are neurotoxic and induce apoptosis ^30,55,56^. Ceramides are the simplest of the sphingolipids and are formed either by *de novo* synthesis from sphingosine or by hydrolysis of sphingomyelin molecules ^57^. While ceramides are rarely found in bacteria, many bacteria in the *Bacteroidetes* phylum can synthesize sphingolipids, which were shown to be taken up and processed by human epithelial cells *in vitro*^58^. Thus, we asked whether variation in ceramide and sphingosine derivatives levels in blood was significantly explained by microbial genera, host genetic factors, or a combination of both.

We observed a large degree of heterogeneity in variance partitioning in ceramide and sphingosine molecules (Fig. 6). Variance in sphingosine itself was only weakly explained by the composition of the gut microbiome (R^2^=0.01), with alpha-diversity as the only significant microbiome-related association. However, other intermediates in ceramide synthesis, such as sphinganine derivatives, showed stronger microbiome and genome associations (i.e. up to 5% of variance explained; Fig. 6A). Ceramides showed a broad range of explained variances. Whereas most ceramides were associated with the gut microbiome, a small subset had additional genetic components to their variances, such as lactosyl-N-nervonoyl-sphingosine and ceramide (d16:1/24:1, d18:1/22:1), both of which had more than 5% of their variation explained by genetic factors. Both of these lipid species highlighted above are examples of very long-chain fatty-acyl sphingolipids. While shorter fatty acids (≤C18) are prevalent in the human diet ^59^, often serving as preferential substrates for ceramide synthesis by some gut microbes ^60^, longer chain fatty acids, such as nervonic acid (24:1), are often the product of elongation by host enzymes before being incorporated into sphingolipids, and are found primarily in brain tissue^61^. In summary, we observed broad heterogeneity in the fraction of variance explained by the microbiome and host genetics, where a difference in the fatty acid chain length could shift a ceramide from being solely associated with microbial genera (i.e. shorter chains) to those mostly associated with host genetics (i.e. longer chains).

**Figure 6.**
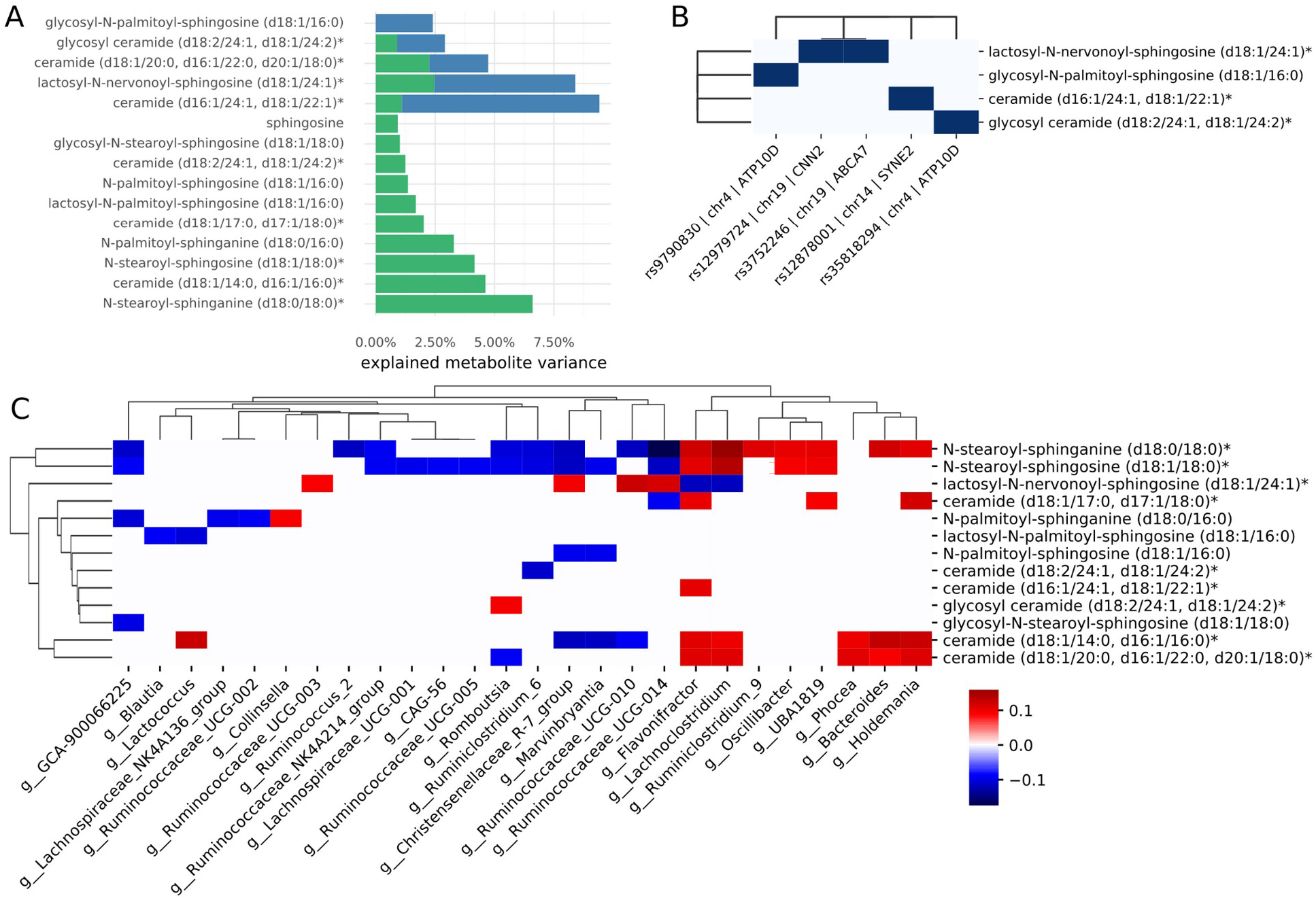
Host genetic and gut microbial associations with sphingosine and ceramides. (A) Explained variances for sphingosine, sphingosine intermediates, and ceramides. (B) Genetic variants associated with sphingosines and ceramides. Dark blue cells denote associations that passed a genome-wide significance threshold in the GWAS. (C) Associations between bacterial genera and sphingosines/ceramides. Only significant associations are shown (FDR-corrected and confounder-adjusted q<0.05). Fill colors denote the correlation coefficients (see legend). Associations were corrected for sex, age, age2, sex:age, and sex:age2 interactions, BMI, microbiome vendor, metabolomics batch, and the first 5 principal components of genetic ancestry. * indicates compounds for which a standard is not available, but Metabolon is confident in its identity; ** indicates a compound for which a standard is not available, but Metabolon is confident in its identity.

## Discussion

In this study, we partition the cross-sectional variance explained in individual blood metabolite levels into their host genetic and gut microbiome components, across a population of 2,049 generally healthy individuals. Prior studies have established that the plasma metabolome is intimately connected to host genetics as well as to the gut microbiome, and that overall genetic and microbial influences on the metabolome appear to be orthogonal. However, it has been unclear as to whether or not genetics and the microbiome act on mutually exclusive sets of metabolites or act simultaneously on individual metabolites. Most of the detected blood metabolites (522/948) showed significant associations with the gut microbiome, which is consistent with prior work. Hybrid genome-microbiome contributions were common and affected about 30% of all metabolites associated with either host genetics or the microbiome. We observed that 3 in 4 metabolites with a significant association with host genetics had additional associations with the gut microbiome. Thus, the majority of blood metabolites associated with host genetics include significant hybrid associations with the gut microbiome. Unlike the set of metabolites associated with host genetics, only 1 in 3 metabolites associated with the microbiome showed an additional hybrid genetic component to their variance. Thus, while both genetic and microbiome variation are important to explaining variation in the blood metabolome, the microbiome (and the myriad factors correlated with variation in the microbiome, like diet and lifestyle) appears to be the dominant driving force.

Additionally, we found some evidence for gene-microbiome interactions, where a particular metabolite-microbiome association was modulated by the genetic background of the host. In general, those interactions only explained small fractions of metabolite variance (<2%). However, this may be a consequence of the generally low prevalence of the minor alleles for the affected genetic variants. These interactions often appear to result from relatively strong associations within the minor allele group, but we have limited numbers of these minor allele carriers to assess these associations robustly. For instance, for homoarginine, we observed a strong negative correlation with *Ruminicocacceae* specific to the heterozygous minor allele background, where low levels or absence of *Ruminicocacceae* in a minority of the heterozygous minor allele population was associated with homoarginine levels close to what was observed in the homozygous major allele carriers. Thus, genetically-determined deviations from a particular quantitative trait may be modulated or even induced by a particular microbial community composition in the gut. These kinds of genome-microbiome interaction effects could help guide the design of microbiome-targeted therapeutics that mitigate host genetic disease risk.

Prior studies of microbe-host co-metabolites derived from microbial precursors in blood, like hippurate or p-cresol sulfate (i.e., hepatically modified forms of microbially-derived metabolites), hinted that the cross-sectional variance in these molecules might only be associated with the microbiome and not with host genetics ^17,38,62,63^. Here we show that those observations do not extrapolate to all microbe-host co-metabolites. Indeed, we found that much of the crosssectional variation in many metabolites derived from gut microbial precursors was explained by host genetics. For example, while unconjugated secondary bile acids in the bloodstream are only associated with the gut microbiome, their glucuronidated or sulfated derivatives, formed in the liver, were strongly influenced by host genetic variation. Most of the variance in deoxycholate glucuronide across the current study cohort could be explained by a combination of genetic and microbial factors, while unconjugated deoxycholate was solely associated with the microbiome. Deoxycholate and deoxycholate glucuronide both showed associations with the microbiome, but these associations were with different sets of bacterial genera. Glucuronidation facilitates excretion into urine and feces and prior work in rats has shown that glucuronidated bile acids are reabsorbed less efficiently than unmodified secondary bile acids in the intestine ^64^. The gut metagenome encodes a variety of β-glucuronidases ^64–66^ which likely enable gut commensals to use these bile-acid-conjugated-glucuronides as a carbon source. Conversely, free secondary bile acids can be toxic to many bacterial taxa ^67^. Thus, bile acids can directly drive changes in the gut microbiome, either by acting as carbon sources that promote growth or as toxic compounds that inhibit growth.

Another intriguing finding from our analyses was the variable association of certain plasma sphingolipids with either host genetics or the gut microbiome, depending on the fatty-acyl groups comprising each lipid species. Ceramides with a ≤C18 fatty-acyl group showed stronger correspondence to the gut microbiome, consistent with the high prevalence of these fatty acids in the diet ^59^ and their preferential incorporation into ceramides by certain gut taxa ^60^. On the other hand, ceramides with very long chain fatty-acyl groups (22:1,24:1), that are most abundant in brain tissue and often synthesized through elongation by host enzymes ^61^, showed a stronger correspondence with host genetics. Importantly, ceramides with different fatty-acyl chain lengths have been implicated in a number of human diseases, including Alzheimer’s disease, depression and mood disorders ^68,69^. Distinguishing which ceramides are under the control of diet and the gut microbiome versus genetic predisposition may aid in the design of new and improved precision therapeutics.

It should be noted that the current study only included individuals from the Pacific West of the U.S. who were predominantly of European descent. While our results are consistent with results obtained from cohorts in Israel and Sweden ^1,70^, future studies in more diverse populations will be required to see whether or not the reported observations replicate more broadly. Finally, while we included many highly relevant covariates in our regression analyses (i.e. age, sex, BMI, and genetic ancestry), many of the observed microbe-metabolite associations are likely confounded with lifestyle and dietary habits, which were not comprehensively tracked in the current study population and can strongly influence the composition of the gut microbiome.

Overall, our analyses show that the plasma metabolome is influenced by a mixture of genetic and microbial factors, where the abundance of individual microbially-derived metabolites absorbed in the gut is often affected additively by both host genetic variation and by variation in the ecology of the gut microbiome. Furthermore, many microbe-metabolite associations are dependent upon the host genetic background. These hybrid genome-microbiome regression models provide unique insights into the forces underlying variation in the human blood metabolome and can suggest possible therapeutic strategies. Finally, we suggest that disease relevant blood metabolites strongly associated with the microbiome may be modifiable through dietary, probiotic, prebiotic or lifestyle interventions, whereas metabolites under genetic control may require pharmacological interventions that target host metabolic pathways. Understanding which of these circulating small molecules fall predominantly under host versus microbiome control will help to guide interventions designed to prevent and/or treat a range of diseases.

## Materials and Methods

### Institutional Review Board Approval for the Study

Procedures for this study were reviewed and approved by the Western Institutional Review Board with the Institutional Review Board (IRB) study number 20170658 for the Institute for Systems Biology and 1178906 for Arivale.

### Cohort Description

All study participants were subscribers of the Arivale Scientific Wellness program and provided consent and research authorization allowing the use of their anonymized, de-identified data in research. The Arivale program is described in detail in Zubair et. al. ^71^. In brief, participants signed up for a comprehensive deep phenotyping program coupled with personalized data-driven wellness coaching in order to improve overall health and wellness. Baseline blood draws were taken at the first in-house visit and paired with at-home fecal sampling. Metabolomics and microbiome measurements taken from the biofluids were described previously in Wilmanski et. al. ^8^ A subset of the participants opted for longitudinal sampling.

### Genome-wide association analysis

DNA from the 2,049 participants was extracted from whole blood samples by Covance (Redmond, WA) using a standardized protocol. Whole genome sequencing was performed by Wuxi, Inc. (Shanghai, China) in a CLIA-certified laboratory. Extracted libraries were sequenced using an HiSeq X sequencer (Illumina, San Diego, USA) with 150bp libraries and aiming for >30x coverage. Basecalling and conversion to raw FASTQ files was performed using the Illumina Basespace software. Raw reads were aligned to the hg19 human reference genome with BWA 0.7.12. The GATK HaplotypeCaller with GATK 3.3.0 was used to call individual variants. This included indel local realignment followed by base quality recalibration. This yielded a set of around 7.68M measured SNPs.

To account for genetic structure and potential cryptic relatedness in the cohort, we used a linear mixed model to test for associations between genetic variants and metabolite levels. For each metabolite, we performed a genome-wide association study using FastGWA using the hg19 genetic map ^72^. Metabolome genome-wide significance was called using the Bonferroni corrected p-value threshold of 5.29e-11 (5e-8 / #metabolites) where 5e-8 is the commonly used threshold for genome-wide significance ^73^. Linkage disequilibrium scores were taken from the 1000 Genomes project ^74^ and used as additional covariate in the genome-wide association study as well as in gene associations calculated with MAGMA ^32^.

### Gut Microbiome Sequencing

Fecal samples were collected using a proprietary at-home swap kit (OMNIgene Gut, DNAGenotek, USA) to maintain DNA conservation and longevity. 250μL of homogenized stool from each sample were subsequently used for DNA extraction using a MoBio PowerMag Soil DNA isolation kit (QIAGEN, Germany) using the KingFisher Flex instrument. DNA concentrations in each sample were quantified with a QuBit (ThermoFisher, USA) and purity was assessed by measuring the A260/A280 absorbance ratio. Amplification and library preparation were performed by external providers using custom and optimized protocols by either amplifying the V4 region (SecondGenome, USA) or the V3-V4 region (DNAGenotek, USA) of the 16S gene.

Sequencing was performed using a MiSeq (Illumina, USA) using either a paired-end 250bp protocol (SecondGenome) or a paired-end 300bp protocol (DNAGenotek). Basecalling was performed using the Illumina Basespace platform which removed the added phiX reads and provided the final FASTQ files used for downstream analysis. Quality of the sequencing reads was assessed by manual inspection of the error rate across sequencing cycles and appropriate length cutoffs of 250bp for the forward reads and 230bp for the reverse reads were chosen based on the profiles. Reads with more than 2 expected errors under the Illumina error model were removed from the analysis along with reads containing ambiguous base calls (“N” nucleotides). More than 97% of reads passed those filters yielding a mean of around 200,000 reads per sample.

Filtered and truncated reads were then used to infer amplicon sequence variants using DADA2 ^75^. Here error profiles were learned for each sequencing run separately. The resulting ASVs and respective counts were merged and chimeras were removed using the “consensus” strategy implemented in DADA2 which removed around 16% of all reads. Taxonomy assignment of ASVs was performed by using the naive Bayes classifier in DADA2 with the SILVA database (version 128). Species assignments were performed by exact match of the inferred ASVs with the reference 16S gene in SILVA where possible. About 90% of reads could be classified down to the genus levels and 32% of reads could be classified down to the species level. The total set of samples was then filtered for those individuals that also had metabolomics data and/or whole genome shotgun sequencing available. Where several microbiome time points were available we used the one closest to the blood draw used for metabolomics. Low abundance taxa, with less than 10 reads on average or appearing in less than 50% of all samples, were removed from the data set.

### Metabolite-microbiome associations and confounder adjustment

Genus-level read counts were scaled using the centered log-ratio (CLR) transform and standardized across the 1,163 samples. Metabolite abundances were log-transformed and standardized which yielded good similarity to a normal distribution as judged by a QQ-plot.

Transformed and scaled metabolite and bacterial genera abundances were then regressed against the set of confounders and the residuals were saved as new dependent variables. For the confounder adjustment we chose the most common intrinsic factors that would affect blood metabolite and fecal microbe abundances. Firstly, we corrected for sex, age, and BMI by including coefficients for biological sex, age, age^2^, sex:age, and sex:age^2^ interactions, and BMI. Additionally, we included covariates for common batch effects like the season of the year the samples were obtained, the metabolomics batch (each sample in a batch was processed together), and the vendor for the microbiome sample (DNA Genotek or Second Genome). Finally, we also included a continuous measure of genetic ancestry given by the first 5 principal components from an analysis of 100,000 linkage disequilibrium corrected frequent SNPs (minor allele frequency > 5%) calculated with the PC-AiR and PC-Relate methods ^76,77^. The residuals were used for all further analyses as well as visualizations and analyses of variances. After this we performed individual linear regressions between all metabolite-genus pairs to find significant associations between individual metabolites and bacterial genera. P-values were obtained using F-tests and corrected for false discovery rate using the method of Benjamini and Hochberg ^78^. We then retained the significant associations with a FDR cutoff q<0.05. We also performed the same analysis with alpha-diversity as the independent variable or with alphadiversity as a covariate to the CLR-transformed bacterial genus abundance to verify that individual genus-level associations were not due to an association with alpha-diversity alone, which we did not observe to be the case.

### Analysis of explained variances (R^2^)

Explained variances (R^2^) for each metabolite were obtained by ordinary least squares models only containing features that were significantly associated with each metabolite individually. Ergo, we only include genus abundances for those genera with a FDR-corrected q<0.05 in the previous metabolite-microbe associations and only those single nucleotide variants that were identified as significant in the previous genome-wide association study for the particular metabolite. This was done to avoid inflation of the explained variances, as adding random variables to a regression model tends to increase the explained variance. We performed multilinear regressions with only microbiome features, only genetic features, and both genetic and microbiome features to quantify the variance of the transformed and confounder-adjusted metabolite abundances that was explained by each feature type individually and jointly. Consequently, it should be noted that the explained variances in this manuscript refer to the metabolite abundance variance after removing components of the variance explained by the covariates listed above. As such, these R^2^ caclulations pose an upper bound of the explained variance of the raw metabolite abundance but are also independent of common confounders, such as age, sex, and BMI and thus should be more generalizable to other populations structures.

Overlap between microbiome and genetic features was quantified as the difference between the sum of R^2^ of the individual feature type models and the joint model R^2^ (R^2^[microbiome] + R^2^[genetics] - R^2^[joint]). In the case of complete independence of microbiome factors and genetics, this difference would be zero and would become positive if there is partial or complete overlap in the variances explained by the microbiome and host genetics.

### Genome-microbiome interactions

For gene microbiome interactions we performed ordinary least squares regression of the form

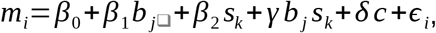

where m_i_ denotes the scaled abundance of metabolite i, b_j_ the scaled abundance of the bacterial genus j, s_k_ the ordinal versions of the allele on variant k (0 - major allele, 1 - heterozygous minor allele, 2 - homozygous major allele), c the vector of confounder covariates, and ε_i_ a random normally distributed variable with an expectation of zero. Significance of the interaction was then evaluated by an F-test comparing the full model to a model with a fixed ɣ=0. P-values from all tested metabolite / variant / genus triplets were adjusted for false discovery rate using the method of Benjamini and Hochberg and judged as significant with a FDR cutoff q<0.05. Even though individual metabolite and bacterial genus abundances had been adjusted for confounders previously, we still included them in this regression because the product bj*sk had not been adjusted for the same confounders. Also, to make our analysis computationally feasible, we performed the regressions only for those metabolites, bacterial genera, and genetic variants that had shown at least one significant interaction in the prior analyses.

Finally, post-hoc tests for microbe-metabolite interactions within sub-cohorts carrying a specific allele were performed by Pearson tests on the product-moment correlation within each subcohort and for those triplets that had shown significant interaction effects.

### Data and Code Availability

Qualified researchers can access the full Arivale deidentified dataset supporting the findings in this study for research purposes through signing a Data Use Agreement (DUA). Inquiries to access the data can be made at and will be responded to within 7 business days. Jupyer notebooks and Python code for reproducing the regression models and figures has been deposited at https://github.com/Gibbons-Lab/2021_gene_environment_interactions.

## Acknowledgments

S.M.G. and C.D. were supported by the Washington Research Foundation Distinguished Investigator Award and startup funds from the Institute for Systems Biology. The funders had no role in designing, carrying out, or interpreting the work presented in the manuscript. T.W. was supported by a generous gift from C. Ellison. Further support came from the National Institutes of Health (NIH) grant (no. U19AG023122) awarded by the National Institute on Aging (NIA) (to N.R.).

## Author contributions

Conceptualization: CLD, CD, SMG, ATM

Methodology: CD, CLD

Formal Analysis: CD, CLD, TW, NR, BS

Investigation: CD, CLD, BS, SMG, ATM

Data Curation: CLD, BS, ATM

Writing - Original Draft: CD, CLD, SMG

Writing - Review & Editing: CD, CLD, TW, PB, SMG, ATM

Visualization: CD, CLD

Supervision & Project Administration: LH, ATM, SMG

---

**Figure S1.**
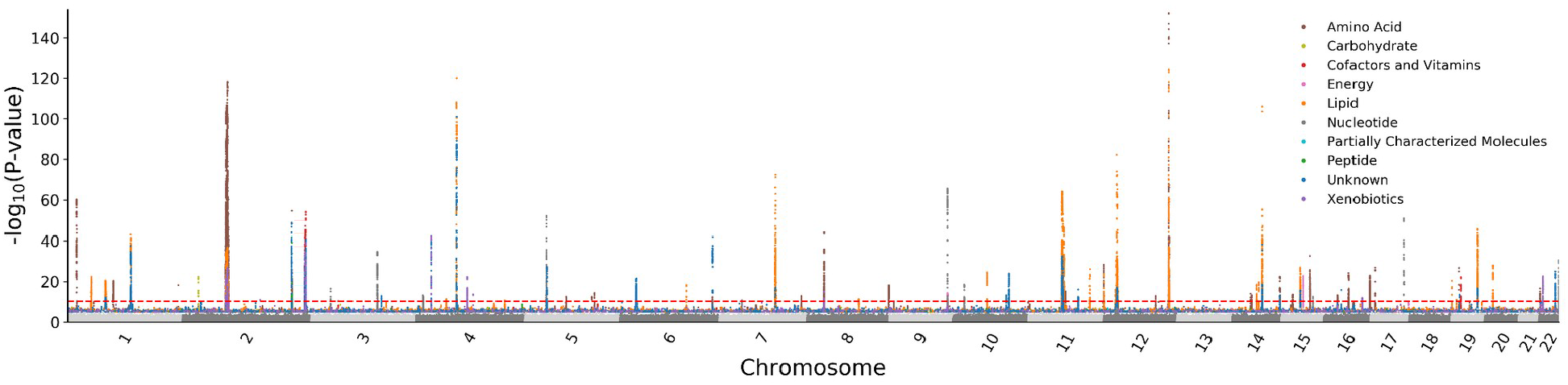
Genome-wide association p-values for variants associated with at least one metabolite. Red dashed line denotes genome-wide significance (p < 5.29e-11).

## Supplementary Data

**Table S1.** R^2^ values for genetics and microbiome for the 594 metabolites with a significant association.

## References

1. Bar, N. et al. A reference map of potential determinants for the human serum metabolome. Nature 588, 135–140 (2020).

2. Wang, X. & Paigen, B. Genetics of variation in HDL cholesterol in humans and mice. Circ. Res. 96, 27–42 (2005).

3. Young, S. G. & Fielding, C. J. The ABCs of cholesterol efflux. Nature genetics vol. 22 316–318 (1999).

4. Blau, N., van Spronsen, F. J. & Levy, H. L. Phenylketonuria. Lancet 376, 1417–1427 (2010).

5. Gieger, C. et al. Genetics meets metabolomics: a genome-wide association study of metabolite profiles in human serum. PLoS Genet. 4, e1000282 (2008).

6. Yu, B. et al. Genetic determinants influencing human serum metabolome among African Americans. PLoS Genet. 10, e1004212 (2014).

7. Gallois, A. et al. A comprehensive study of metabolite genetics reveals strong pleiotropy and heterogeneity across time and context. Nat. Commun. 10, 4788 (2019).

8. Wilmanski, T. et al. Blood metabolome predicts gut microbiome α-diversity in humans. Nat. Biotechnol. 37, 1217–1228 (2019).

9. Rothschild, D. et al. Environment dominates over host genetics in shaping human gut microbiota. Nature 555, 210–215 (2018).

10. Goodrich, J. K. et al. Human genetics shape the gut microbiome. Cell 159, 789–799 (2014).

11. Goodrich, J. K. et al. Genetic Determinants of the Gut Microbiome in UK Twins. Cell Host Microbe 19, 731–743 (2016).

12. Nicholson, J. K. et al. Host-Gut Microbiota Metabolic Interactions. Science 336, 1262–1267 (2012).

13. Forman, B. M. et al. Identification of a nuclear receptor that is activated by farnesol metabolites. Cell 81, 687–693 (1995).

14. Jiao, Y., Lu, Y. & Li, X.-Y. Farnesoid X receptor: a master regulator of hepatic triglyceride and glucose homeostasis. Acta Pharmacol. Sin. 36, 44–50 (2015).

15. Zheng, X. et al. Hyocholic acid species improve glucose homeostasis through a distinct TGR5 and FXR signaling mechanism. Cell Metab. 33, 791–803.e7 (2021).

16. Kawamata, Y. et al. A G protein-coupled receptor responsive to bile acids. J. Biol. Chem. 278, 9435–9440 (2003).

17. Lees, H. J., Swann, J. R., Wilson, I. D., Nicholson, J. K. & Holmes, E. Hippurate: the natural history of a mammalian-microbial cometabolite. J. Proteome Res. 12, 1527–1546 (2013).

18. Nakamura, A. et al. Symbiotic polyamine metabolism regulates epithelial proliferation and macrophage differentiation in the colon. Nat. Commun. 12, 2105 (2021).

19. Tofalo, R., Cocchi, S. & Suzzi, G. Polyamines and Gut Microbiota. Front Nutr 6, 16 (2019).

20. Ohtani, N. & Kawada, N. Role of the gut-liver axis in liver inflammation, fibrosis, and cancer: A special focus on the gut Microbiota relationship. Hepatol. Commun. 3, 456–470 (2019).

21. Vitali, C., Khetarpal, S. A. & Rader, D. J. HDL Cholesterol Metabolism and the Risk of CHD: New Insights from Human Genetics. Curr. Cardiol. Rep. 19, 132 (2017).

22. Moschetta, A., Bookout, A. L. & Mangelsdorf, D. J. Prevention of cholesterol gallstone disease by FXR agonists in a mouse model. Nat. Med. 10, 1352–1358 (2004).

23. Kenny, D. J. et al. Cholesterol Metabolism by Uncultured Human Gut Bacteria Influences Host Cholesterol Level. Cell Host Microbe 28, 245–257.e6 (2020).

24. Dayama, G., Priya, S., Niccum, D. E., Khoruts, A. & Blekhman, R. Interactions between the gut microbiome and host gene regulation in cystic fibrosis. Genome Med. 12, 12 (2020).

25. Hunter, D. J. Gene-environment interactions in human diseases. Nat. Rev. Genet. 6, 287–298 (2005).

26. Naj, A. C. et al. Common variants at MS4A4/MS4A6E, CD2AP, CD33 and EPHA1 are associated with late-onset Alzheimer’s disease. Nat. Genet. 43, 436–441 (2011).

27. Liu, G. et al. Analyzing large-scale samples confirms the association between the ABCA7 rs3764650 polymorphism and Alzheimer’s disease susceptibility. Mol. Neurobiol. 50, 757–764 (2014).

28. Cuyvers, E. et al. Mutations in ABCA7 in a Belgian cohort of Alzheimer’s disease patients: a targeted resequencing study. Lancet Neurol. 14, 814–822 (2015).

29. Cutler, R. G. et al. Involvement of oxidative stress-induced abnormalities in ceramide and cholesterol metabolism in brain aging and Alzheimer’s disease. Proc. Natl. Acad. Sci. U. S. A. 101, 2070–2075 (2004).

30. Mielke, M. M. et al. Serum ceramides increase the risk of Alzheimer disease: the Women’s Health and Aging Study II. Neurology 79, 633–641 (2012).

31. Huynh, K. et al. Concordant peripheral lipidome signatures in two large clinical studies of Alzheimer’s disease. Nat. Commun. 11, 5698 (2020).

32. de Leeuw, C. A., Mooij, J. M., Heskes, T. & Posthuma, D. MAGMA: generalized gene-set analysis of GWAS data. PLoS Comput. Biol. 11, e1004219 (2015).

33. Bosma, P. J. et al. Bilirubin UDP-glucuronosyltransferase 1 is the only relevant bilirubin glucuronidating isoform in man. J. Biol. Chem. 269, 17960–17964 (1994).

34. Gaibar, M., Novillo, A., Romero-Lorca, A., Esteban, M. E. & Fernández-Santander, A. Pharmacogenetics of ugt genes in North African populations. Pharmacogenomics J. 18, 609–612 (2018).

35. Dupuis, J. et al. New genetic loci implicated in fasting glucose homeostasis and their impact on type 2 diabetes risk. Nat. Genet. 42, 105–116 (2010).

36. Wikoff, W. R. et al. Metabolomics analysis reveals large effects of gut microflora on mammalian blood metabolites. Proc. Natl. Acad. Sci. U. S. A. 106, 3698–3703 (2009).

37. Teufel, R. et al. Bacterial phenylalanine and phenylacetate catabolic pathway revealed. Proc. Natl. Acad. Sci. U. S. A. 107, 14390–14395 (2010).

38. Pallister, T. et al. Hippurate as a metabolomic marker of gut microbiome diversity: Modulation by diet and relationship to metabolic syndrome. Sci. Rep. 7, 13670 (2017).

39. Brial, F. et al. Human and preclinical studies of the host-gut microbiome co-metabolite hippurate as a marker and mediator of metabolic health. Gut 70, 2105–2114 (2021).

40. Ridlon, J. M., Kang, D. J., Hylemon, P. B. & Bajaj, J. S. Bile acids and the gut microbiome. Curr. Opin. Gastroenterol. 30, 332–338 (2014).

41. Winham, S. J. & Biernacka, J. M. Gene-environment interactions in genome-wide association studies: current approaches and new directions. J. Child Psychol. Psychiatry 54, 1120–1134 (2013).

42. März, W. et al. Homoarginine, cardiovascular risk, and mortality. Circulation 122, 967–975 (2010).

43. Pilz, S. et al. Low homoarginine concentration is a novel risk factor for heart disease. Heart 97, 1222–1227 (2011).

44. de Aguiar Vallim, T. Q., Tarling, E. J. & Edwards, P. A. Pleiotropic roles of bile acids in metabolism. Cell Metab. 17, 657–669 (2013).

45. King, C. D., Rios, G. R., Green, M. D. & Tephly, T. R. UDP-glucuronosyltransferases. Curr. Drug Metab. 1, 143–161 (2000).

46. Alnouti, Y. Bile Acid sulfation: a pathway of bile acid elimination and detoxification. Toxicol. Sci. 108, 225–246 (2009).

47. Xiang, X. et al. Effect of SLCO1B1 polymorphism on the plasma concentrations of bile acids and bile acid synthesis marker in humans. Pharmacogenet. Genomics 19, 447–457 (2009).

48. Uhlén, M. et al. Tissue-based map of the human proteome. Science 347, 1260419 (2015).

49. Tóth, B. et al. Human OATP1B1 (SLCO1B1) transports sulfated bile acids and bile salts with particular efficiency. Toxicol. In Vitro 52, 189–194 (2018).

50. Meikle, P. J. & Summers, S. A. Sphingolipids and phospholipids in insulin resistance and related metabolic disorders. Nat. Rev. Endocrinol. 13, 79–91 (2017).

51. Chaurasia, B. et al. Targeting a ceramide double bond improves insulin resistance and hepatic steatosis. Science 365, 386–392 (2019).

52. Norris, G. H., Porter, C. M., Jiang, C., Millar, C. L. & Blesso, C. N. Dietary sphingomyelin attenuates hepatic steatosis and adipose tissue inflammation in high-fat-diet-induced obese mice. J. Nutr. Biochem. 40, 36–43 (2017).

53. Xia, J. Y. et al. Targeted Induction of Ceramide Degradation Leads to Improved Systemic Metabolism and Reduced Hepatic Steatosis. Cell Metab. 22, 266–278 (2015).

54. Grassi, S. et al. Thematic Review Series: Biology of Lipid Rafts: Lipid rafts and neurodegeneration: structural and functional roles in physiologic aging and neurodegenerative diseases. J. Lipid Res. 61, 636 (2020).

55. Toledo, J. B. et al. Metabolic network failures in Alzheimer’s disease: A biochemical road map. Alzheimers. Dement. 13, 965–984 (2017).

56. Green, D. R. Apoptosis and sphingomyelin hydrolysis. The flip side. The Journal of cell biology vol. 150 F5–7 (2000).

57. Summers, S. A., Chaurasia, B. & Holland, W. L. Metabolic messengers: Ceramides. Nat Metab 1, 1051–1058 (2019).

58. Johnson, E. L. et al. Sphingolipids produced by gut bacteria enter host metabolic pathways impacting ceramide levels. Nat. Commun. 11, 2471 (2020).

59. Ervin, R. B., Wright, J. D., Wang, C.-Y. & Kennedy-Stephenson, J. Dietary intake of fats and fatty acids for the United States population: 1999-2000. Adv. Data 1–6 (2004).

60. Stankeviciute, G. et al. Convergent evolution of bacterial ceramide synthesis. Nat. Chem. Biol. 1–8 (2021).

61. Fan, Y., Meng, H.-M., Hu, G.-R. & Li, F.-L. Biosynthesis of nervonic acid and perspectives for its production by microalgae and other microorganisms. Appl. Microbiol. Biotechnol. 102, 3027–3035 (2018).

62. Clayton, T. A., Baker, D., Lindon, J. C., Everett, J. R. & Nicholson, J. K. Pharmacometabonomic identification of a significant host-microbiome metabolic interaction affecting human drug metabolism. Proc. Natl. Acad. Sci. U. S. A. 106, 14728–14733 (2009).

63. Tevzadze, G. et al. Effects of a gut microbiome toxin, p-cresol, on the indices of social behavior in rats. Neurophysiology 50, 372–377 (2018).

64. Oelberg, D. G., Little, J. M., Adcock, E. W. & Lester, R. Intestinal absorption of bile acid glucuronides in rats. Dig. Dis. Sci. 33, 1110–1115 (1988).

65. Creekmore, B. C. et al. Mouse Gut Microbiome-Encoded β-Glucuronidases Identified Using Metagenome Analysis Guided by Protein Structure. mSystems 4, (2019).

66. Gloux, K. et al. A metagenomic β-glucuronidase uncovers a core adaptive function of the human intestinal microbiome. Proc. Natl. Acad. Sci. U. S. A. 108 Suppl 1, 4539–4546 (2011).

67. Begley, M., Gahan, C. G. M. & Hill, C. The interaction between bacteria and bile. FEMS Microbiol. Rev. 29, 625–651 (2005).

68. Dinoff, A., Herrmann, N. & Lanctôt, K. L. Ceramides and depression: A systematic review. J. Affect. Disord. 213, 35–43 (2017).

69. Tong, M. & de la Monte, S. M. Mechanisms of ceramide-mediated neurodegeneration. J. Alzheimers. Dis. 16, 705–714 (2009).

70. Dekkers, K. F. et al. An online atlas of human plasma metabolite signatures of gut microbiome composition. medRxiv 2021.12.23.21268179 (2021) doi:10.1101/2021.12.23.21268179.

71. Zubair, N. et al. Genetic Predisposition Impacts Clinical Changes in a Lifestyle Coaching Program. Sci. Rep. 9, 6805 (2019).

72. Jiang, L. et al. A resource-efficient tool for mixed model association analysis of large-scale data. Nat. Genet. 51, 1749–1755 (2019).

73. Xu, C. et al. Estimating genome-wide significance for whole-genome sequencing studies. Genet. Epidemiol. 38, 281–290 (2014).

74. 1000 Genomes Project Consortium et al. A global reference for human genetic variation. Nature 526, 68–74 (2015).

75. Callahan, B. J. et al. DADA2: High-resolution sample inference from Illumina amplicon data. Nat. Methods 13, 581–583 (2016).

76. Conomos, M. P., Miller, M. B. & Thornton, T. A. Robust inference of population structure for ancestry prediction and correction of stratification in the presence of relatedness. Genet. Epidemiol. 39, 276–293 (2015).

77. Conomos, M. P., Reiner, A. P., Weir, B. S. & Thornton, T. A. Model-free Estimation of Recent Genetic Relatedness. Am. J. Hum. Genet. 98, 127–148 (2016).

78. Benjamini, Y. & Hochberg, Y. Controlling the False Discovery Rate: A Practical and Powerful Approach to Multiple Testing. Journal of the Royal Statistical Society: Series B (Methodological) 57, 289–300 (1995).

